# ERK1/2 Inhibition Alleviates Noise-Induced Hearing Loss While Tempering Down the Immune Response

**DOI:** 10.1101/2023.10.18.563007

**Authors:** Richard D. Lutze, Matthew A. Ingersoll, Alena Thotam, Anjali Joseph, Joshua Fernandes, Tal Teitz

## Abstract

Noise-induced hearing loss (NIHL) is a major cause of hearing impairment, yet no FDA-approved drugs exist to prevent it. Targeting the mitogen activated protein kinase (MAPK) cellular pathway has emerged as a promising approach to attenuate NIHL. Tizaterkib is an orally bioavailable, highly specific ERK1/2 inhibitor, currently in Phase-1 anticancer clinical trials. Here, we tested tizaterkib’s efficacy against permanent NIHL in mice at doses equivalent to what humans are currently prescribed in clinical trials. The drug given orally 24 hours after noise exposure, protected an average of 20-25 dB SPL in three frequencies, in female and male mice, had a therapeutic window >50, and did not confer additional protection to KSR1 genetic knockout mice, showing the drug works through the MAPK pathway. Tizaterkib shielded from noise-induced cochlear synaptopathy, and a 3-day, twice daily, treatment with the drug was the optimal determined regimen. Importantly, tizaterkib was shown to decrease the number of CD45 and CD68 positive immune cells in the cochlea following noise exposure, which could be part of the protective mechanism of MAPK inhibition.

## Introduction

Hearing loss afflicts more than 10% of the world population with noise-induced hearing loss (NIHL) as one of the main causes^1–4^. NIHL has increased over the past years due to the use of personal headphones, noisy city environments, and noise exposure in military combat^5–9^. Hearing loss occurs with intense noise exposure due to mechanical damage to stereocilia and cellular mechanisms that lead to cell death and dysfunction^10–13^. Inhibiting pathways that induce cochlear stress and dysfunction is a promising approach to limit the amount of hearing loss that occurs with damaging noise exposures^11,14^. There are currently no Food and Drug Administration (FDA)-approved drugs for the treatment of NIHL; therefore, there is a clinical need to develop compounds that can protect individuals from this very common disorder^4,15,16^.

Recently, our laboratory has identified several inhibitors of the mitogen activated protein kinase (MAPK) pathway as promising compounds that protect from noise- and cisplatin-induced hearing loss^4^. The MAPK pathway is involved in a multitude of cellular processes and is a common target for many diseases^17^. The MAPK pathway is commonly deregulated in various types of cancer but it has been implicated in hearing loss over the last 15-20 years^4,18–21^. This cellular pathway consists of a phosphorylation cascade of different proteins that are activated when phosphorylated. Protein ERK1/2 is the main kinase in this cascade and activates downstream pathways and transcription factors^22^. Activation of the MAPK pathway has traditionally been associated with cell proliferation and survival but this is dependent on the cell type and stimulus^17,22^. Cells in the inner ear are post-mitotic and not actively proliferating and several studies have demonstrated that activation of this pathway in these types of tissues can cause cellular stress, dysfunction, and ultimately, cell death^4,19,23–26^. Due to these findings in post-mitotic cochlear cells, targeting ERK1/2 is a promising approach to mitigate NIHL.

Tizaterkib (formerly known as AZD-0364) is a newly developed, orally bioavailable, highly specific ERK1/2 inhibitor that is currently in Phase-1 clinical trials for the treatment of advanced solid tumors and hematological malignancies^27,28^. It was demonstrated to have a low IC_50_ in vitro of 6 nM and a high affinity for the ERK1/2 kinases^28^. Additionally, our laboratory has shown that tizaterkib protects cochlear explants from cisplatin-induced outer hair cell (OHC) death with an IC_50_ of 5 nM, close to the IC_50_ value of 6 nM measured for inhibition of the ERK1/2 kinase activity in cell lines^4,28^. Due to the high specificity of the drug to inhibit ERK1/2 and low predicted doses that will limit off-target effects, tizaterkib is a very useful tool to study ERK1/2’s role in NIHL^27,28^. Targeting ERK directly in the MAPK pathway may be more efficient for protection from hearing loss because ERK modulates different downstream pathways that have been implicated in NIHL such as cellular death and inflammation^22,26,29,30^. Additionally, our laboratory demonstrated that dabrafenib, a BRAF kinase inhibitor, protects mice from NIHL. The use of an ERK inhibitor would confirm that inhibition of the MAPK pathway, not just the upstream molecular target BRAF, mitigates NIHL^4^.

One cellular mechanism that is associated with NIHL is the induction of the inflammatory response which produces an immune response in the cochlea^11,31–35^. Several studies have shown that inhibition of cytokines and chemokines protects mice from NIHL and other types of hearing loss^36–40^. Additionally, dexamethasone, a corticosteroid which acts as an anti-inflammatory agent, has been shown to be protective against many types of hearing loss, including NIHL^41,42^. Furthermore, infiltrating immune cells from the peripheral blood have been implicated in the damage to cochlear cells as a result of a bystander effect in which immune cells release cytokines and chemokines which exacerbates that inflammatory response and can cause cellular dysfunction and eventually cell death^32,35,43,44^. This suggests that the inflammatory response which leads to an overactive immune response in the cochlea is partially responsible for the damage that occurs after noise insults^45^. The MAPK pathway has been shown to regulate the inflammatory and immune response and one of the possible ways that MAPK inhibition protects from NIHL is through moderating this critical cellular response following noise^23,46–50^.

In this study, we perform a dose-response of tizaterkib in an animal model to determine the minimum effective oral dose that protects from NIHL. We examine whether ERK1/2 inhibition mitigates noise-induced synaptopathy which commonly occurs with damaging noise exposures, and demonstrate that tizaterkib protects from NIHL primarily through the MAPK pathway by employing the KSR1 KO genetic mouse model^51,52^. We also show that tizaterkib mitigates NIHL not only after 100-dB SPL, 2 hours’ noise exposure, but also at a higher noise exposure levels of 106-dB SPL for 2 hours, which widens the therapeutic potential of the drug. Furthermore, we performed different schedules of administration of the drug to determine the optimal treatment regimen to protect from NIHL with MAPK inhibition. Finally, we test whether ERK1/2 inhibition alters the immune response following noise exposure to elucidate one of tizaterkib’s mechanisms of protection from NIHL.

## Results

### Tizaterkib protects from noise-induced hearing loss when treatment begins 45 minutes before noise exposure

We first determined whether tizaterkib (chemical structure in Figure 1A) protects from NIHL when mice were pretreated with the drug. Briefly, mice were orally treated with 25 mg/kg/bw tizaterkib, a dose previously published with no side effects to mice^27,28^, 45 minutes before noise exposure. Mice were exposed to 100 dB SPL noise for 2 hours at the 8-16 kHz octave band. This noise exposure induces permanent threshold shifts in FVB/NJ mice^4,15^. Mice were then orally treated again in the evening and treatment proceeded for another 2 days, for a total of 3 days of treatment, twice a day (Figure 1B). Mice treated with tizaterkib had significantly lower Auditory Brainstem Response (ABR) threshold shifts at the 8, 16, and 32 kHz frequencies compared to the noise alone cohort. Tizaterkib treated mice had an average reduction of ABR threshold shifts of 27 dB at 8 kHz, 16 dB at 16 kHz, and 15 dB at 32 kHz compared to noise alone mice (Figure 1C). Tizaterkib alone treated mice had no hearing loss which indicates the drug causes no ototoxicity when administered by itself. In addition, the mice treated with 25 mg/kg/bw twice a day for three days did not display any behavioral changes or signs of general toxicity as determined by weight measurements compared to carrier alone treated mice (Figure 1D).

**Figure 1:**
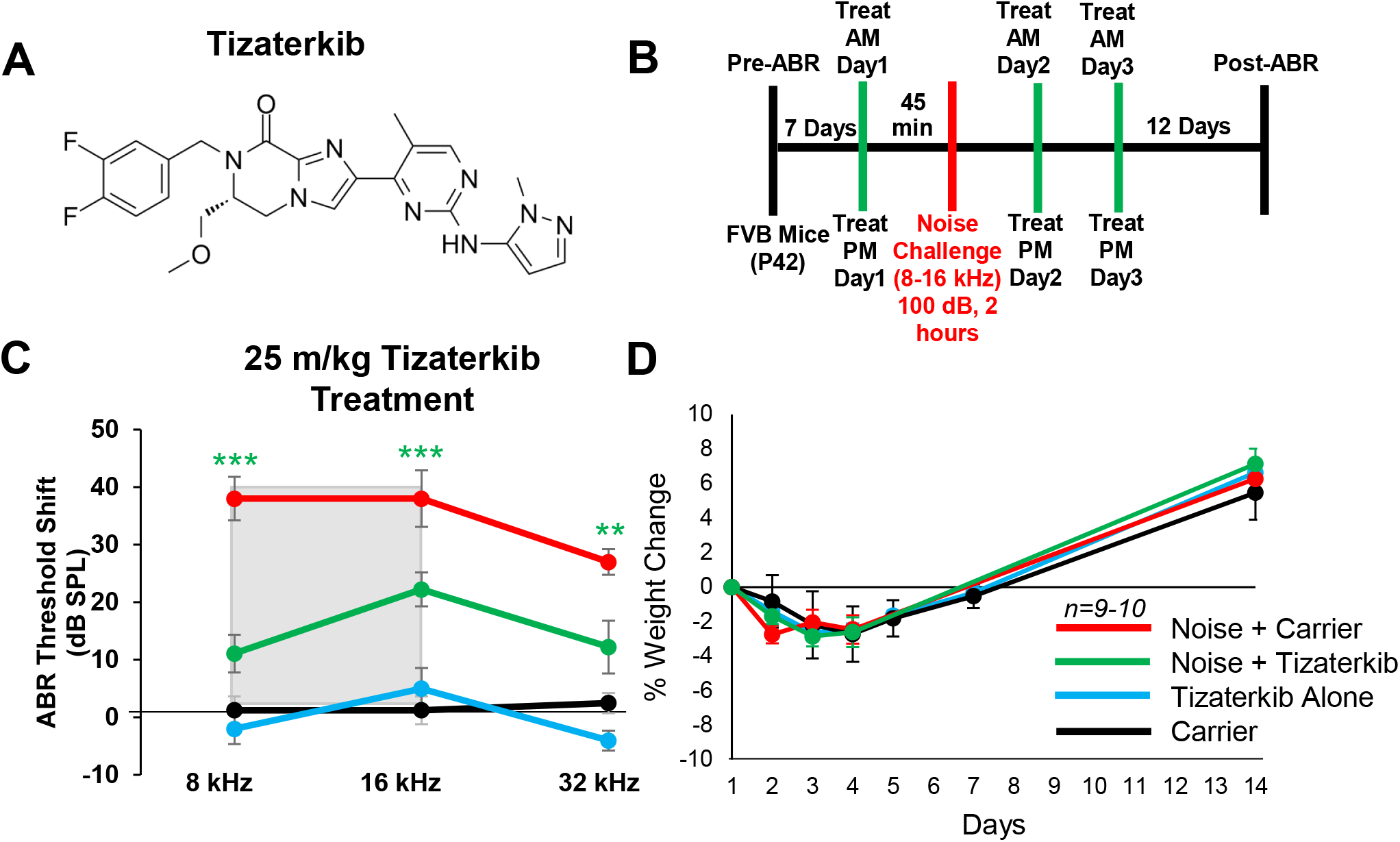
AZD0364 protects mice from noise-induced hearing loss when administered 45 minutes before noise exposure. **(A)** Molecular structure of tizaterkib. **(B)** Schedule of administration for noise exposure and tizaterkib treatment in FVB mice. Mice were given first treatment of tizaterkib via oral gavage 45 minutes before noise exposure. Mice were treated with the drug for a total of 3 days, twice a day and given noise exposure once. **(C)** ABR threshold shifts following procedure in (B). **(D)** Percent weight change of different experimental cohorts throughout the 14-day protocol shown in (B). Noise + Carrier (red), noise + tizaterkib (green), tizaterkib alone (blue), and carrier (black). Data shown as means ± SEM, *P<0.05, **P<0.01, ***P<0.001 compared to noise alone by two-way ANOVA with Bonferroni post hoc test. *n=9-10* mice

### Tizaterkib administration protects from NIHL when treatment starts 24 hours after noise exposure

Noise exposure cannot always be predicted; therefore, we determined if protection still occurs when treatment begins after noise exposure. Figure 2A shows the treatment and noise exposure protocol in which tizaterkib treatment began 24 hours after noise exposure and mice were orally treated twice a day for three total days. Post-noise exposure treatment of 25 mg/kg tizaterkib significantly lowered ABR threshold shifts compared to noise alone treated mice. An average threshold shift reduction of 20 dB at 8 kHz, 25 dB at 16 kHz, and 10 dB at 32 kHz was observed compared to the noise alone cohort (Figure 2B). 5 mg/kg tizaterkib conferred the same amount of protection as 25 mg/kg (Figure 2B). A lower dose of 0.5 mg/kg was then tested, which also significantly lowered ABR threshold shifts at the 8 and 16 kHz frequencies. An average threshold shift reduction of 22 dB at 8 kHz and 23 dB at 16 kHz was observed compared to noise alone treated mice (Figure 2C & Expanded View Figure 1). 0.1 mg/kg tizaterkib was then tested to determine the minimum effective dose and 0.1 mg/kg did not offer significant protection, which indicates that 0.5 mg/kg was close to the minimum effective dose to protect from NIHL (Figure 2D). Figure 2E shows a dose response curve of protection from NIHL at the 16 kHz frequency with 25, 5, and 0.5 mg/kg all offering similar levels of protection. Males and females were analyzed separately to test if there were any sex differences and tizaterkib equally protects both sexes from NIHL (Figure 2F & G).

**Figure 2:**
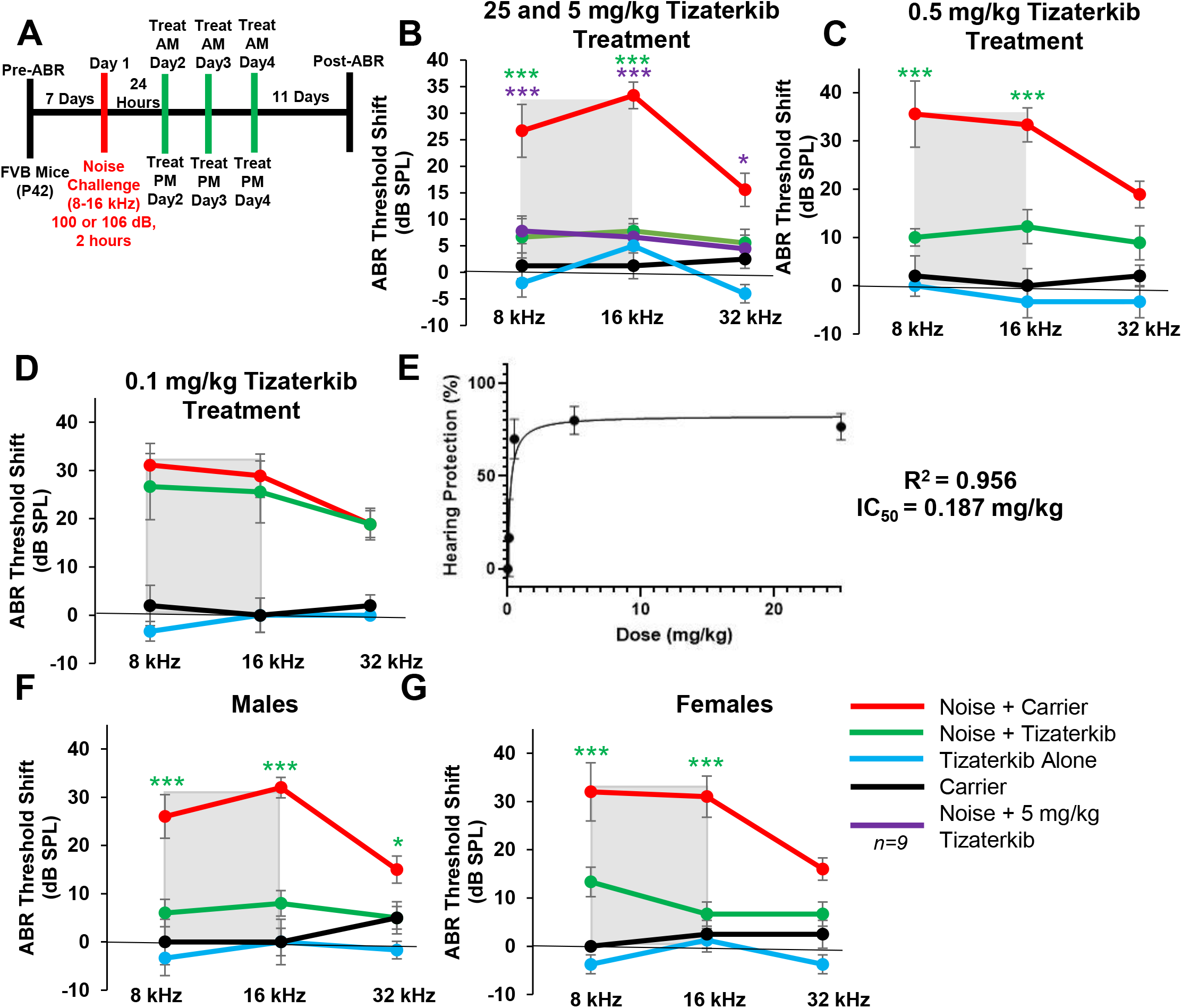
Tizaterkib protects mice from noise-induced hearing loss when administered 24 hours after noise exposure. **(A)** Schedule of noise exposure and tizaterkib which began 24 hours after noise exposure. Mice were treated with varying concentrations of the tizaterkib twice a day for 3 total days. **(B)** ABR threshold shifts following procedure in (A) with 25 and 5 mg/kg tizaterkib given to separate groups. **(C)** ABR threshold shifts following procedure in (A) with 0.5 mg/kg administered to mice. **(D)** ABR threshold shifts following procedure in (A) with 0.1 mg/kg tizaterkib treatment. **(E)** Dose-response curve of tizaterkib protection from noise induced hearing loss at 16 kHz with 100% protection as a 0 dB SPL threshold shift. **(F)** ABR threshold shifts of males treated with tizaterkib following procedure in (A). **(G)** ABR threshold shifts of females treated with tizaterkib following procedure in (A). Noise + carrier (red), noise + tizaterkib (green), noise + 5mg/kg tizaterkib (purple), tizaterkib alone (blue), and carrier (black). Data shown as means ± SEM, *P<0.05, **P<0.01, ***P<0.001 compared to noise alone by two-way ANOVA with Bonferroni post hoc test. *n= 9-10* mice

### Tizaterkib protects from noise-induced cochlear synaptopathy

Following the post treatment hearing tests, mouse cochleae were collected and stained with Ctbp2 and myosin VI to determine whether ERK1/2 inhibition protects from cochlear synaptopathy, which occurs following noise exposure^4,15,53^. Representative images of cochlear samples stained with Ctbp2 and myosin VI are shown in Figure 3A & C, which represent the 8 and 16 kHz regions, respectively. The number of ctbp2 puncta per inner hair cell (IHC) were quantified for the 8 and 16 kHz region (Figures 3B & D). The average number of ctbp2 puncta per IHC at the 8 and 16 kHz region in non-noise exposed mice was 12.3 and 14.2, respectively. The noise alone cohort had an average of 9.1 and 9.7 ctpb2 puncta per IHC at the 8 and 16 kHz regions, respectively. 0.5 mg/kg tizaterkib treated mice had significantly more ctbp2 puncta compared to noise alone mice with 11.9 at 8 kHz and 11.5 at 16 kHz (Figure 3 B & D). Additionally, tizaterkib treated mice had significantly higher ABR wave 1 amplitude at 16 kHz following noise exposure compared to noise alone mice at 90 dB SPL (Figure 3E).

**Figure 3:**
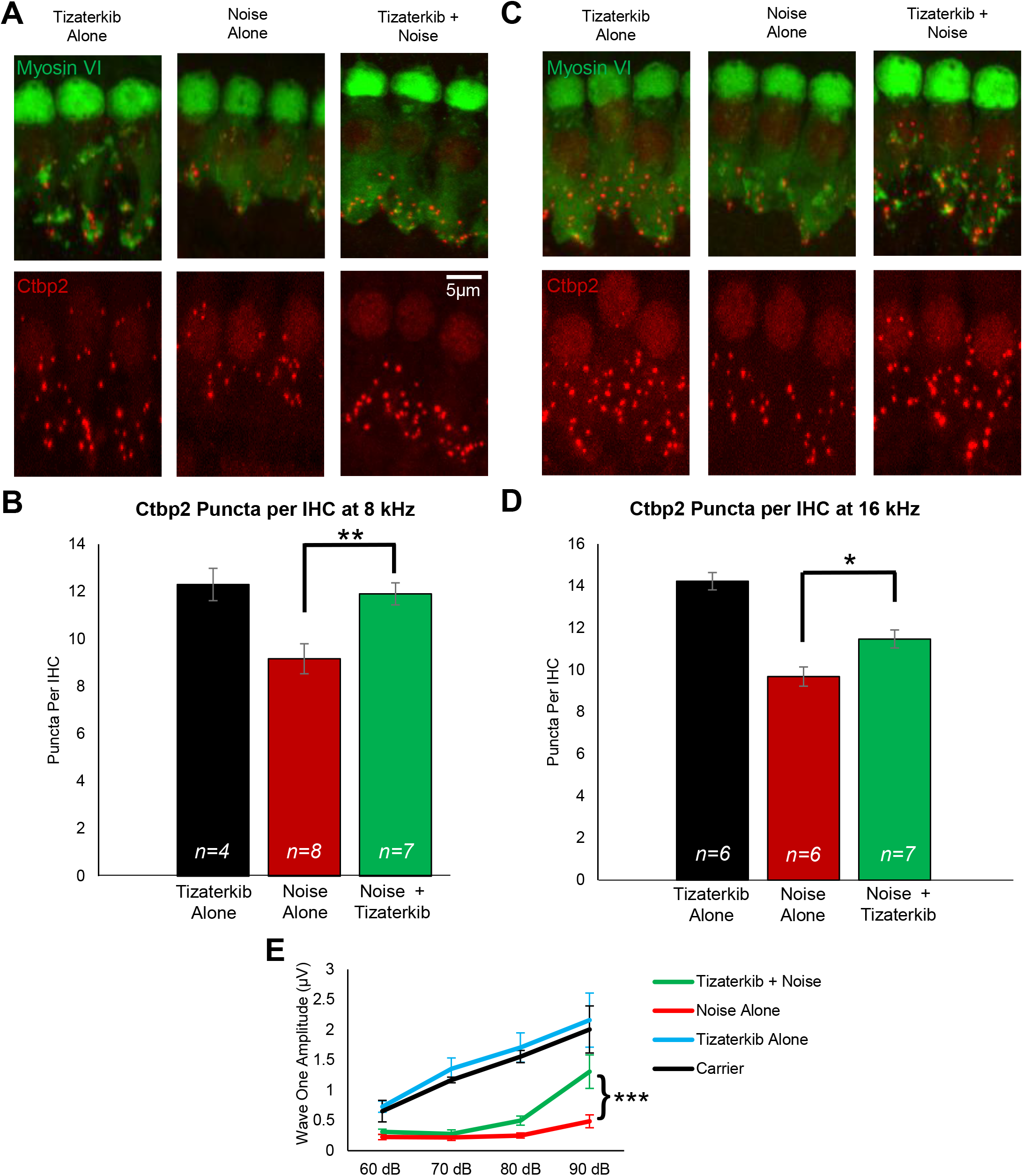
Tizaterkib protects mice from noise-induced synaptopathy at the 8 and 16 kHz regions. **(A)** Representative images of whole mount cochlear sections stained with myosin VI (green) and Ctbp2 (red) at the 8 kHz region. **(B)** Number of Ctbp2 puncta per IHC at the 8 kHz region. **(C)** Representative images of whole mount cochlear sections at the 16 kHz region. **(D)** Number of Ctbp2 puncta per IHC at the 16 kHz region. Data shown as means ± SEM, *P<0.05, **P<0.01 compared to noise alone by one-way ANOVA with Bonferroni post hoc test. Tizaterkib Alone (black), Noise Alone (red), Noise + Tizaterkib (green).

### Tizaterkib protects mice from NIHL when exposed to a higher noise exposure intensity of 106 dB SPL

In this experiment, the same drug treatment protocol as before was followed, except mice were exposed to 106 dB SPL noise instead of 100 dB SPL (Figure 4A). The noise alone group had in average 10 dB higher ABR threshold shifts compared to mice exposed to 100 dB (Figure 2). Mice orally treated with 0.5 mg/kg tizaterkib had significantly lower ABR threshold shifts at the 8 and 16 kHz regions (Figure 4B). Tizaterkib treated mice had lower average threshold shifts of 15 dB at 8 kHz and 22 dB at 16 kHz compared to noise alone mice. Additionally, tizaterkib treated mice following noise exposure have significantly lower DPOAE threshold shifts at 16 kHz with an average reduction of 14 dB SPL compared to noise alone mice (Figure 4C).

**Figure 4:**
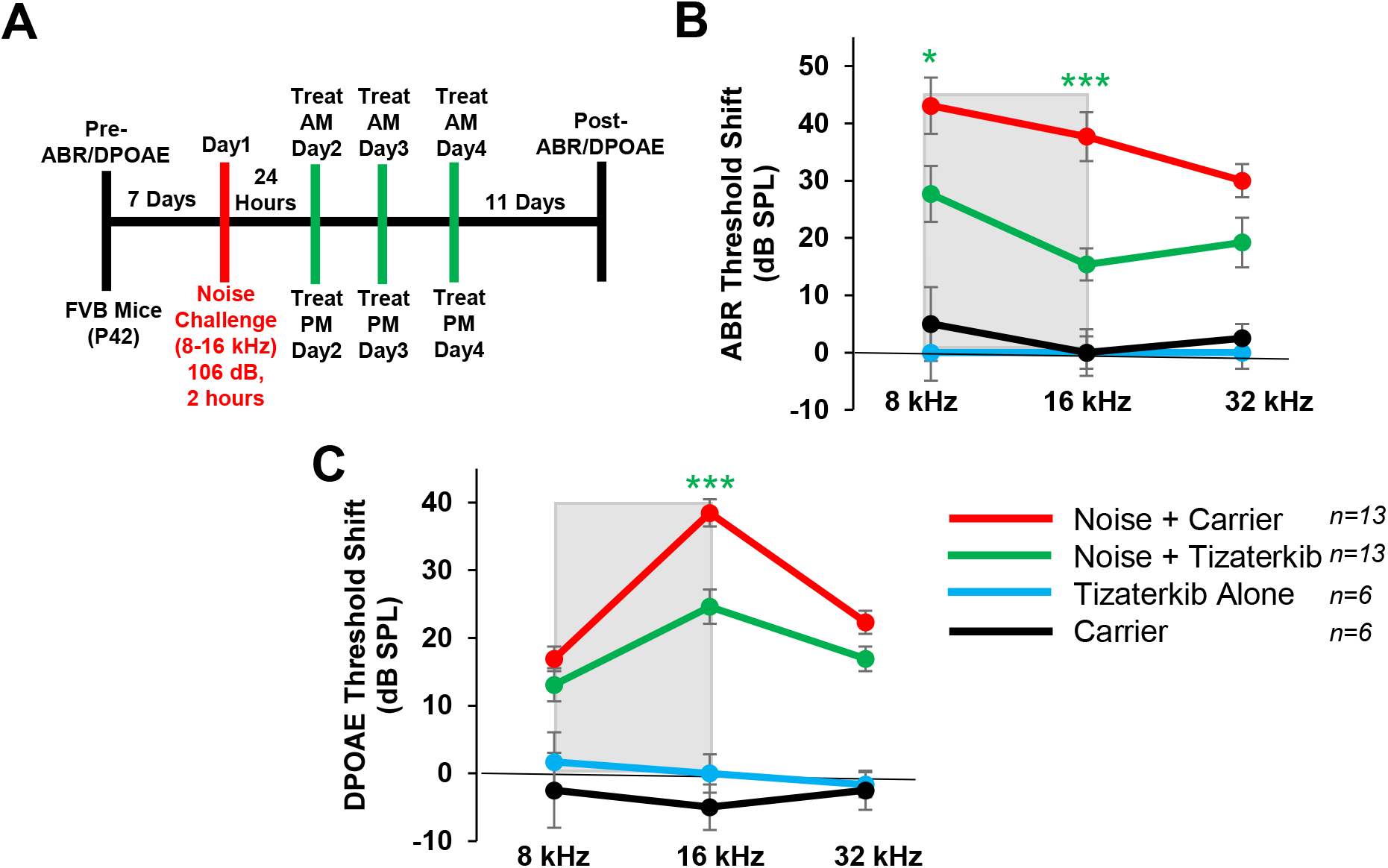
Tizaterkib protects from noise induced hearing loss when mice are exposed to 106 dB. **(A)** Schedule of noise exposure and tizaterkib treatment. Mice were exposed to 106 dB SPL for 2 hours and tizaterkib treatment started 24 hours after noise exposure. Mice were treated for 3 total days, twice a day. **(B)** ABR threshold shifts following protocol in (A). **(C)** DPOAE threshold shifts following protocol in (A). Noise + Carrier (red), noise + tizaterkib (green), tizaterkib alone (blue), and carrier (black). Data shown as means ± SEM, *P<0.05, **P<0.01, ***P<0.001 compared to noise alone by two-way ANOVA with Bonferroni post hoc test. *n= 13* mice

### Tizaterkib treatment in KSR1 KO mice does no confer any extra protection from NIHL

Tizaterkib was expected to protect mice from NIHL through ERK inhibition and this was tested by employing KSR1 genetic germline knockout (KO) and wild type (WT) mouse littermates. KSR1 is a scaffolding protein in the MAPK pathway and eliminating KSR1 protein significantly lowers MAPK activation and activity (Figure 5A)^51,52^. KSR1 WT and KO littermate mice, C57BL/6 strain, were exposed to 100 dB SPL, 8-16 kHz octave band, noise for 2 hours and then treated by oral gavage with tizaterkib or carrier twice a day for three days, starting 24 hours post noise exposure (Figure 5B). Tizaterkib treatment in KSR1 WT mice offers significant protection from NIHL at 16 and 32 kHz with an average reduction in ABR threshold shifts of 18 and 19 dB, respectively (Figure 5C). KO mice alone and KO mice treated with tizaterkib have almost identical protection from NIHL with average reductions in threshold shifts of 27 dB at 16 kHz and 28 dB at 32 kHz. There is no significant difference in threshold shifts between the WT mice treated with tizaterkib and either KO cohort exposed to noise (Figure 5C).

**Figure 5:**
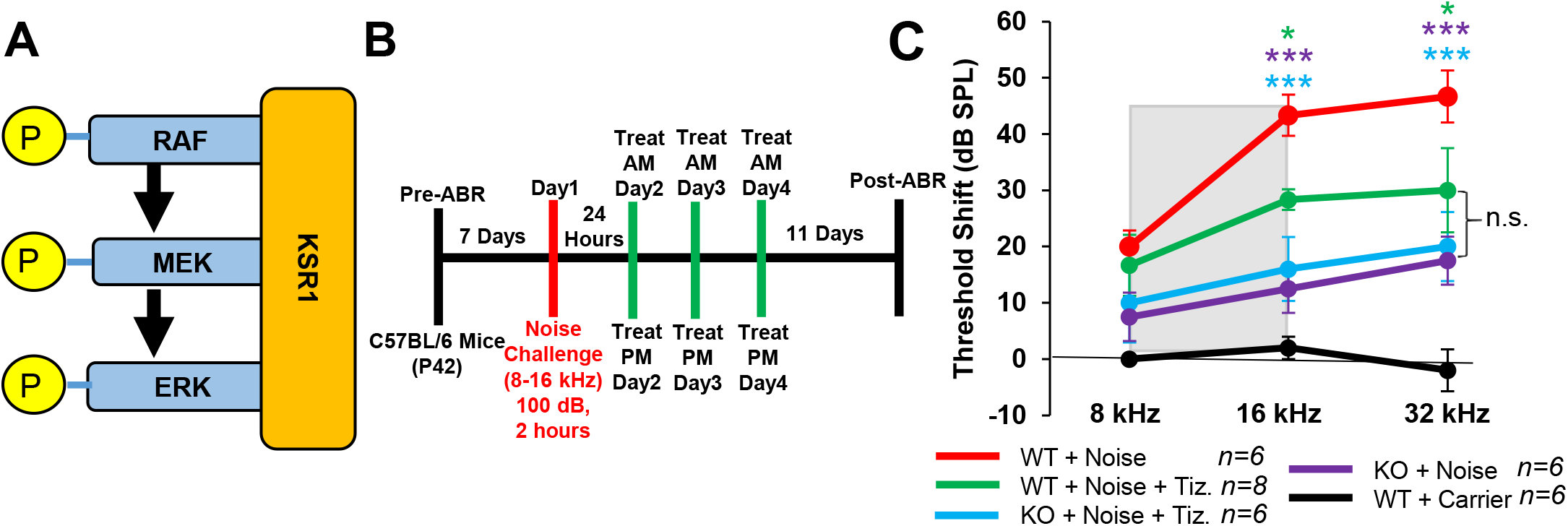
Tizaterkib treatment phenocopies the resistance to noise-induced hearing loss measured in the KSR1 KO mouse model. **(A)** KSR1 is a scaffolding protein for RAF, MEK, and ERK which enables efficient transmission of MAPK signals. **(B)** Schedule of noise exposure and 5 mg/kg tizaterkib treatment in KSR1 WT and KO mice. Mice were exposed to 100 dB SPL for 2 hours and tizaterkib treatment began 24 hours after noise exposure. Mice were treated with tizaterkib or carrier twice a day for 3 total days. **(C)** ABR threshold shifts following protocol in (B). WT + noise (red), KO + noise + tizaterkib (blue), WT + noise + tizaterkib (green), KO + noise (purple), and WT + carrier alone (black). Data shown as means ± SEM, *P<0.05, **P<0.01, ***P<0.001 compared to noise alone by two-way ANOVA with Bonferroni post hoc test. *n=5-6* mice

### Three Days of oral Tizaterkib treatment produces better protection from NIHL compared to 1 and 2 days of treatment

Mice treated with 0.5 mg/kg tizaterkib twice a day for three days offers significant protection from NIHL when treatment begins 24 hours after noise exposure. We checked if a delay in the first treatment to 48 hours following noise exposure would still offer significant protection. We performed the same noise and tizaterkib treatment protocol except the first tizaterkib treatment occurred 48 hours after noise and not 24 hours (Figure 6A). Mice treated with 0.5 mg/kg tizaterkib did not have significantly lower ABR threshold shifts compared to noise alone mice, even though they did trend lower (Figure 6B). We then determined whether a single day or 2 days of treatment starting 24 hours after noise instead of 3 days offers similar protection (Figure 6C). Mice treated with 0.5 mg/kg tizaterkib for one day did have significantly lower threshold shifts at 32 kHz and mice treated with tizaterkib for 2 days have lower threshold shifts at 8 and 16 kHz with slightly better protection than 1 day of treatment (Figure 6D). Three days of treatment was significantly better than 1 and 2 days of treatment with an average reduction in threshold shifts of 12, 15, and 11 dB at the 8, 16, and 32-kHz regions, respectively, compared to mice treated for 2 days (Figure 6D).

**Figure 6:**
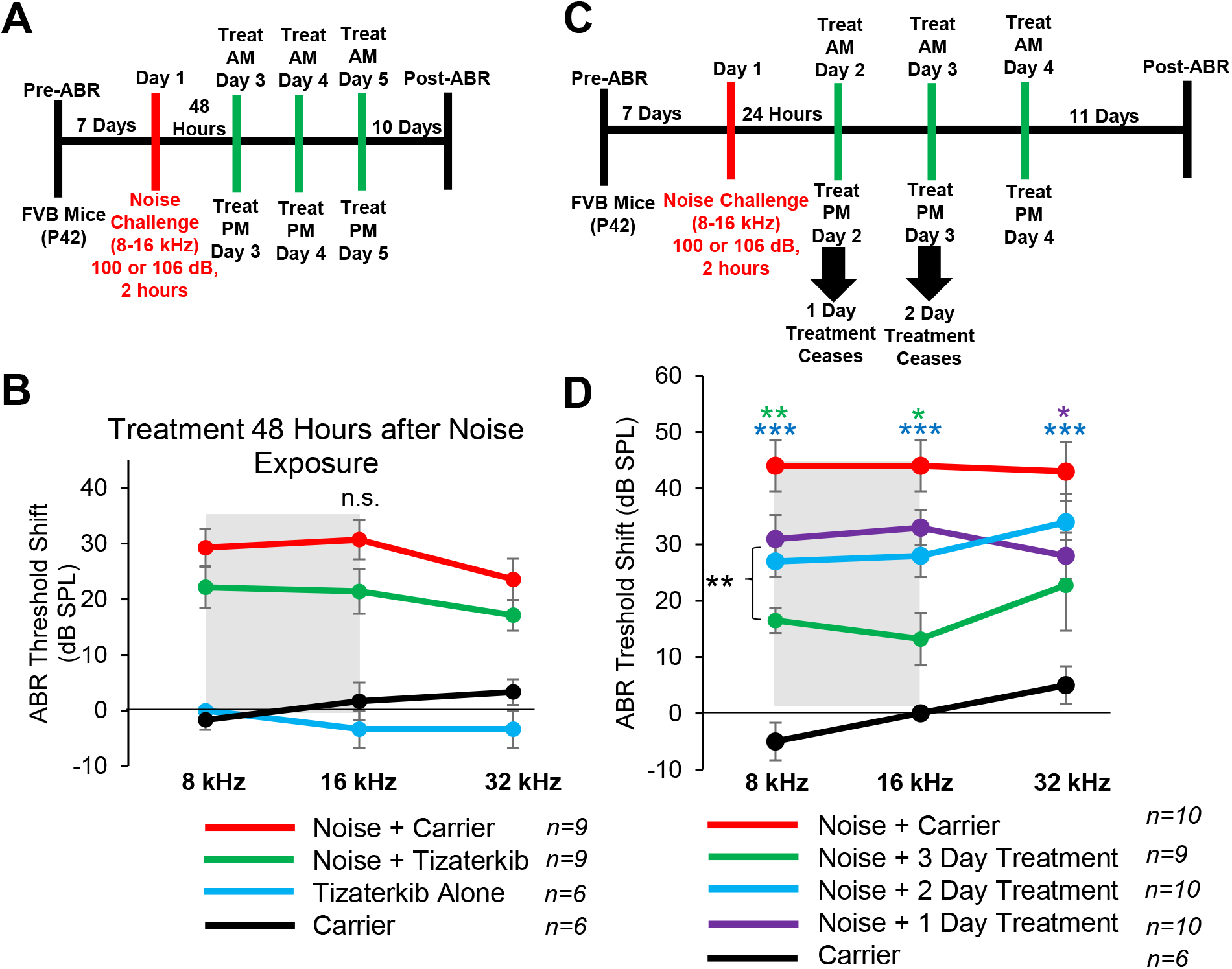
3 Days of treatment beginning 24 hours after noise exposure produces optimal protection with tizaterkib administration. **(A)** Schedule of administration for noise exposure and tizaterkib treatment. Treatment with tizaterkib began 48 hours after noise exposure and mice were treated for 3 days, twice a day. **(B)** ABR threshold shifts following the protocol in (A). noise alone (red), noise + tizaterkib (green), carrier (black), and tizaterkib alone (blue). **(C)** Schedule of administration for noise exposure and tizaterkib treatment. Treatment began 24 hours after noise exposure and one cohort was treated for 1 day, one cohort was treated for 2 days, and another cohort was treated for 3 days. **(D)** ABR threshold shifts following the protocol in (C). noise alone (red), 1-day treatment + noise (purple), 2-day treatment + noise (blue), 3-day treatment + noise (green), carrier alone (black). Data shown as Data shown as means ± SEM, *P<0.05, **P<0.01, ***P<0.001 compared to noise alone by two-way ANOVA with Bonferroni post hoc test.

### Tizaterkib treated mice have significantly less immune cells in their cochleae compared to noise alone treated mice

Modulating the number of immune cells in the cochlea following noise insult could lead to protection form NIHL^54–56^. We exposed the mice to the same noise exposure as before (100 dB for 2 hours at 8-16 kHz octave band) and treated mice with carrier or 0.5 mg/kg tizaterkib for three days, twice a day. Mice were then sacrificed 1 hour after the last drug treatment and stained with anti-CD45 antibody to determine the number of total immune cells in the cochlea following noise insult. Mice exposed to noise and treated with carrier had a significant increase in the number of CD45 positive cells in their cochleae and tizaterkib treatment significantly lowered the number of CD45 positive cells compared to noise alone to almost carrier or drug alone levels (Figure 7 A & C). The scala tympani region was also analyzed further because this region had the largest increase in immune cells following noise insult. Again, the density of CD45+ cells in the scala tympani region was increased in noise alone mice, was significantly lower in tizaterkib treated mice exposed to noise, and reached the levels of the carrier or drug alone mice (Figure 7 B & D). There was a 1.8 -fold increase in CD45 positive cells between the noise alone and the noise + tizaterkib treated mice (Figure 7C) and an even larger fold-increase of 2.4 in CD45 positive cells in the scala tympani region between the noise and noise + tizaterkib treated mice (Figure 7D). Additionally, there was a 30% decrease in CD45 positive cells in the stria vascularis for tizaterkib treated mice following noise exposure compared to the noise alone cohort (Expanded View Figure 2). Cochlear cryosections stained with secondary antibody alone (no CD45 primary antibody) did not have positive CD45 staining (Expanded View Figure 3A) and normal immune cell morphology was observed when higher magnification images were taken of CD45 positive cells (Expanded View Figure 3B).

**Figure 7:**
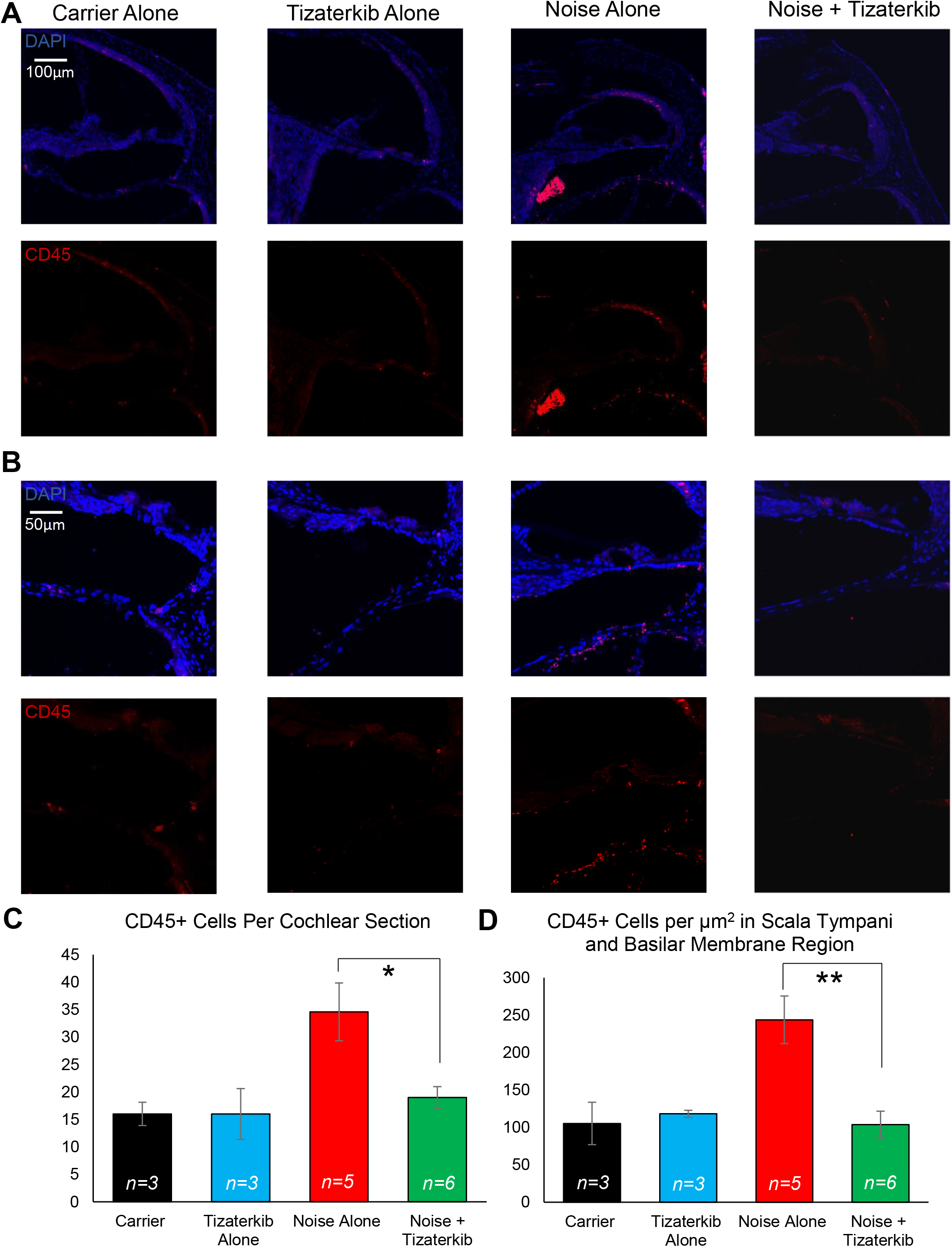
Tizaterkib treatment lowers the number of CD45 positive cells in the cochlea 4 days’ post noise exposure. **(A)** Representative low magnification images of cochlear cryosections stained with CD45 (red) and DAPI (blue). The treatment protocol shown in Figure 2A was utilized and mice were sacrificed 4 days following noise exposure 1 hour after the final tizaterkib treatment. **(B)** Higher magnification of images shown in (A) of the scala tympani and basilar membrane region. **(C)** Quantification of CD45 positive cells of cochlear sections in (A). **(D)** Quantification of CD45 positive cells per µm^2^ in the scala tympani as represented in (B). carrier (black), tizaterkib alone (blue), noise alone (red), noise + tizaterkib (green). Data shown as means ± SEM, *P<0.05, **P<0.01 compared to noise alone by one-way ANOVA with Bonferroni post hoc test. n=3-6

Total cochlear protein lysates were prepared from mice sacrificed 6 days following noise exposure to follow the duration in which this difference in immune cells number persists. Western blots were run and probed with anti-CD45 antibody (general immune cells marker), anti-CD68 antibody (a macrophage marker), and anti-GAPDH antibody (a loading control). There was a 1.6-fold difference in the amount of CD45 in the cochlea on day 6 and a 1.4-fold difference in CD68 between the noise alone and noise + tizaterkib treated mice (Figure 8). When mice were sacrificed 8 days following noise insult instead of 4 or 6 days, and cochlear cryosections were stained with CD45, the number of CD45 positive cells in the cochlea following noise exposure returned to normal levels (Expanded View Figure 4).

**Figure 8:**
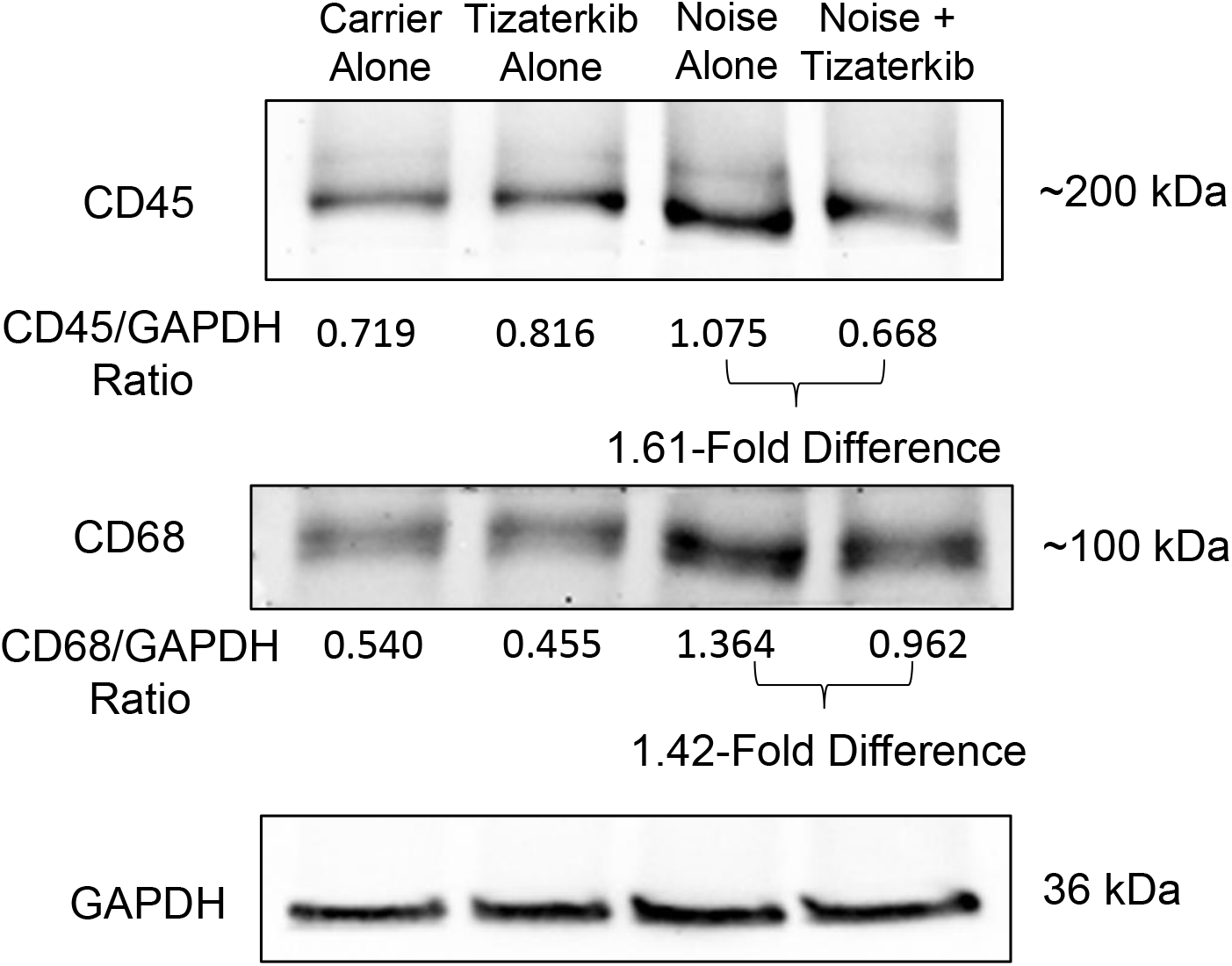
Tizaterkib treatment lowers the amount of CD45 and CD68 in the cochlea 6 days’ post noise exposure. Western blots showing the amount of CD45 and CD68 in the cochlea following noise exposure and tizaterkib treatment. The same treatment protocol shown in Figure 2A was utilized and mice were sacrificed 6 days after noise exposure. The CD45/GAPDH and CD68/GAPDH ratios are shown underneath the western blot. GAPDH was used as the loading control. The experimental groups from left to right are as follows: carrier alone, tizaterkib alone, noise alone, and noise + tizaterkib. Each group had the cochleae from 5 mice (10 cochleae) pooled together to make the tissue lysate.

## Discussion

Tizaterkib mitigates NIHL at a low dose of 0.5 mg/kg/bw when treatment begins 24 hours after noise exposure and mice are treated twice a day for three days. A total of 1 mg/kg/bw is administered per day which is the mouse equivalent to what humans are currently receiving in phase-1 clinical trials for cancer treatment^57,58^. Tizaterkib has a wide therapeutic window of at least 50 in mice for the treatment of NIHL-25 mg/kg offers significant protection with no known deleterious side effects and 0.5 mg/kg offers almost identical protection which makes that therapeutic window at least 50 in mice. A wide therapeutic window is necessary for any drug to get to clinical trials so physicians can have a wide range of doses to give patients with minimal side effects while still getting the desired effect and accounting for individual differences in drug metabolism^14,59–61^. This preliminary data on tizaterkib is promising because it demonstrates that ERK1/2 can be targeted for protection from hearing loss with little to no side effects occurring, which has been a legitimate concern of ERK1/2 inhibitors in the past.

Tizaterkib protects from NIHL when the drug is first administered 45 minutes before noise exposure and treatment continues after noise exposure. The drug offers identical protection when treatment begins 24 hours after noise exposure compared to treatment beginning 45 minutes before the insult. This is encouraging because noise exposure cannot always be predicted so treatment beginning after noise exposure is more translationally relevant and widens the therapeutic applications of targeting the MAPK pathway. Additionally, 3 days of treatment was shown to offer significantly better protection compared to 1 and 2 days of treatment. These treatment timing experiments demonstrate the optimal times to target the MAPK pathway after noise exposure and show critical times to intervene with treatment in order to have protection from NIHL. Translationally, tizaterkib is administered to patients in phase-1 clinical trials for 21 days and we got significant protection from hearing loss with only 3 days of treatment with the same daily dose^57^. This makes tizaterkib a promising preclinical compound for the treatment of NIHL. The total amount of drug that is given to mice is less than what is currently being administered to humans which is very crucial when repurposing drugs for the treatment of noise-induced hearing loss.

ERK1/2 inhibition not only protects from permanent NIHL, but it also protects mice from cochlear synaptopathy, which commonly occurs after damaging noise exposures^4,15,62^. Tizaterkib treated mice have more Ctbp2 puncta per IHC compared to noise alone treated mice which demonstrates that less synaptic dysfunction occurs with ERK1/2 inhibition. Additionally, tizaterkib treated mice have larger ABR wave 1 amplitude, which is a functional correlate of cochlear synaptopathy^63^. This is an important finding for two main reasons. 1) Part of the protective effect that occurs from ERK1/2 inhibition could be through prevention of synaptic dysfunction^4,15^. Tizaterkib treated mice have less synaptic dysfunction and permanent hearing loss compared to noise alone mice, so protection of synapses could be leading to protection from permanent hearing loss. 2) Protection from cochlear synaptopathy also lowers the risk of future age related hearing loss^64–66^. Synaptic dysfunction where the IHC synapses with the auditory nerve fibers has been shown to increase the risk of age related hearing loss and other hearing dysfunctions, such as speech recognition and hearing in a noisy environment^67,68^.

Tizaterkib protected mice from NIHL at a low dose of 0.5 mg/kg when mice were exposed to 100 dB SPL for two hours; therefore, we wanted to test a higher noise exposure level to examine whether ERK inhibition also protects at higher noise levels. Exposing FVB mice to 100 dB for 2 hours induces permanent hearing loss but we wanted to test the drug’s ability to protect from a more intense noise exposure level^4,15^. This study demonstrates that tizaterkib protects mice from NIHL when the animals are exposed to 106 dB SPL for two hours. The level of protection following 106 dB SPL noise exposure is 90% of the 100 dB 2-hour noise protection (Figure 2C & 4B). We also show that DPOAE threshold shifts were lower with tizaterkib treatment which suggests ERK inhibition protects OHCs from noise induced dysfunction. OHCs are one of the main cell types in the inner ear affected by noise exposure so this is important data to demonstrate^11,69–71^. Protection from more intense noise exposures demonstrates the therapeutic advantage of targeting the MAPK pathway for mitigation of NIHL. Previous studies have shown that some drugs protect from 100 dB noise exposure but not 106 dB which makes targeting this pathway even more attractive and promising^15^.

The KSR1 mouse model was utilized to demonstrate that tizaterkib was protecting through inhibition of the MAPK pathway and not through some other nonspecific off target effects. We have recently shown that KSR1 KO mice have reduce phosphorylation of ERK1/2 in their cochleae after noise exposure, and are partially resistant to noise-induced hearing loss compared to their WT KSR1 littermates (Manuscript in Preparation Ingersoll et al.). Here, we show that pharmacological inhibition of molecular target ERK1/2 achieves similar resistance levels to the KSR1 KO mice. If the protective effect of tizaterkib was occurring through an off target effect, we would expect that KSR1 KO mice treated with tizaterkib would have a difference in protection from NIHL compared to KSR1 KO mice not treated with the drug. Both KO mouse groups, KO alone and KO plus tizaterkib, had identical protection to one another and WT KSR1 mice treated with the drug were not significantly different from either KO group. This supports our hypothesis that tizaterkib is protecting mice from NIHL through inhibition of the MAPK pathway, which was expected due to the drug’s specificity for ERK1/2, but nonetheless needed to be tested^27,28^. In the past, a main concern of MAPK inhibitors was their off target effects and toxicity profiles^72^, but this result demonstrates that tizaterkib is not protecting through off target effects but directly through inhibition of ERK1/2 and the MAPK pathway.

Immune cell infiltration to the cochlea following noise exposure has been implicated as a possible mechanism that is leading to hearing loss^34,40,73^. This study demonstrates that ERK1/2 inhibition lowers the number of infiltrating immune cells following noise insult. We used CD45 as a general marker for all immune cells to determine if ERK inhibition affected the entire immune response following noise, not a specific subset of immune cells. Previous studies have shown that the increase in CD45 positive cells is due to infiltrating immune cells and not through proliferation of resident macrophages^43^. We observed a major increase in CD45 positive cells in the walls of the scala tympani, the spiral ligament region, and the stria vascularis. This is in agreement with previous studies that have shown this increase in immune cells in the same regions of the cochlea that we observed^43,73–75^. Limiting the number of immune cells could protect cochlear cells from secondary damage that can occur with the infiltration of CD45 positive cells ^35,40^.

In our study, we first chose to look at the number of CD45 positive cells in the cochlea 4 days following noise exposure because previous reports show that the peak in infiltrating immune cells occurs at this time point^35^. We saw a significant difference between the noise alone mice and mice treated with tizaterkib. Additionally, we still observed more CD45 protein in noise exposed cochlea 6 days after noise and tizaterkib treated mice had lower amounts. We also observed lower amounts of CD68 protein, a macrophage marker, in the cochlea of tizaterkib treated mice compared to noise alone animals. Previous studies have demonstrated that up to 95% of the immune cells in the cochlea are from the monocyte/macrophage origin; therefore, we are proposing that tizaterkib is mainly lowering the number of monocytes that are infiltrating into the cochlea which then differentiate into macrophages^34,35,43,73^. This was supported by the decrease of CD68 protein quantified in the total protein lysates of cochleae of tizaterkib treated mice compared to noise alone mice on day 6 after noise exposure (Figure 8). We had another cohort of mice whose cochlea were analyzed 8 days following noise exposure and the number of CD45 positive cells were back to baseline levels in the noise alone mouse cohort (Extended View Figure 4). This is in agreement with other studies that showed there was no longer an increased number of immune cells in the cochlea 7-8 days after noise exposure^34,35,76,77^.

There are several interesting points raised when observing the immune cell data from this study. 1) ERK1/2 inhibition could be protecting mice from NIHL through lowering the number of infiltrating immune cells into the cochlea. This is a possible mechanism of protection that will further be explored to determine the role of immune cells in NIHL. 2) Our treatment protocol beginning 24 hours after noise and continuing for 3 days’ lines up with the same time that most CD45 positive cells are infiltrating the cochlea. Immune cell infiltration starts to occur between 24 and 48 hours after noise exposure and peaks around the 4-day mark, the same time point that the mice are receiving treatments^35,43,73,76^. This could explain why 3 days of treatment confers better protection from NIHL compared to 1 and 2 days of treatment (Figure 6 & 7). 3) This study further implicates that infiltrating immune cells could be a contributing factor to NIHL. There is a correlation between immune cell infiltration and hearing loss as shown by the present study and others^33,35,40^, but future studies will have to look more in depth to determine whether immune cells number above a specific threshold are a causative factor leading to hearing loss following noise exposure. Additionally, future studies will look at the different immune cell populations to determine which ones are most affected by ERK inhibition following noise exposure. Even though it has been demonstrated that most immune cells in the cochlea are of the monocyte/macrophage origin, neutrophils are another type of immune cell that is increased in the cochlea following noise insult^33,78,79^. Furthermore, we would like to determine exactly how ERK inhibition is lowering the number of immune infiltrates into the cochlea. ERK has been shown to be involved in cellular stress and death pathways and inhibiting these pathways following noise could indirectly lower the number of immune cells^23,24,26^. Inhibiting cellular stress would lower the amount of pro-inflammatory molecules produced, such as cytokines, chemokines, and reactive oxygen species (ROS), which would then lower the number of immune cells infiltrating from the periphery^35,36,39,40^. Activation of ERK can also affect immune cell migration so ERK inhibition could be directly inhibiting the migration of immune cells to the cochlea following noise insult^80,81^. Interesting to note that ERK inhibitors may also play a key role in sensing damage levels and reducing immune response in other post-mitotic cells outside the ear, such as in neurodegenerative disorders of the kidney, joints, and brain^82,83^.

In summary, we show that tizaterkib, a highly specific ERK1/2 inhibitor, protects mice from NIHL at clinically relevant doses and in two mouse strains – FVB/NJ and C57BL/6 (KSR1 mice). The best time to start treatment is 24 hours following noise exposure and treatment continues for a total of 3 days. The drug protects both female and male mice with equal efficiency. This treatment regimen partially protects mice from cochlear synaptopathy, which will likely decrease the risk of future presbycusis, as damage from noise exposure is a common risk factor for age-related hearing loss. Additionally, tizaterkib protects mice from levels of noise exposure of 100 and 106-dB SPL, and the protective mechanism was shown to be through the MAPK pathway and not by other off target pathways. Finally, ERK1/2 inhibition was shown to lower the number of CD45 and CD68 positive immune cells in the inner ear following noise exposure which could be part of the protective mechanism of tizaterkib. This study further supports that targeting the MAPK pathway is a promising therapeutic strategy for mitigating NIHL, and the ERK1/2 inhibitor, tizaterkib, is an intriguing compound that needs to be further studied as a possible drug for alleviating NIHL in humans.

## Materials and Methods

### Ethics Statement

All animal procedures were approved by the Institutional Animal Care and Use Committee of Creighton University (IACUC) in accordance with policies established by the Animal Welfare Act (AWA) and Public Health Service (PHS).

### Mouse Models

FVB/NJ mice were purchased from Jackson Laboratory and used as breeders in the Creighton University animal research facility. All FVB/NJ mice were 6-8 weeks old at the start of each experiment. KSR1 mice on the C57BL/6 background were a kind gift from Dr. Robert Lewis from the University of Nebraska Medical Center, Omaha, NE. KSR1 heterozygous mice were bred to get KSR1 KO and WT littermates. All KSR1 mice used for experiments were 6-7 weeks old at the beginning of the experiment. All mice were cared for by the laboratory and animal facility staff.

### Auditory Brainstem Response

ABR waveforms in anesthetized mice were recorded in a sound booth by using subdermal needles positioned in the skull, below the pinna, and at the base of the tail, and the responses were fed into a low-impedance Medusa digital biological amplifier system (RA4L; TDT; 20-dB gain). Mice were anesthetized by 500 mg/kg Avertin (2,2,2-Tribromoethanal, T4, 840-2; Sigma-Aldrich) with full anesthesia determined via toe pinch. At the tested frequencies (8, 16, and 32 kHz), the stimulus intensity was reduced in 10-dB steps from 90 to 10 dB to determine the hearing threshold. ABR waveforms were averaged in response to 500 tone bursts with the recorded signals filtered by a band-pass filter from 300 Hz to 3 kHz. ABR threshold was determined by the presence of at least 3 of the 5 waveform peaks. Baseline ABR recordings were performed when mice were 7-8 weeks old and post experimental recordings were performed 14 days after noise exposure. All beginning threshold values were between 10 and 40 dB at all tested frequencies. All thresholds were determined independently by two-three experimenters for each mouse who were blind to the treatment the mice received. Threshold shifts were calculated by subtracting the pre-noise exposure recording from the post-noise exposure recording. ABR wave one amplitudes were measured as the difference between the peak of wave 1 and the noise floor of the ABR trace.

### Distortion Product Otoacoustic Emission

Distortion product otoacoustic emissions were recorded in a sound booth while mice were anesthetized. Mice were anesthetized by 500 mg/kg Avertin (2,2,2-Tribromoethanal, T4, 840-2; Sigma-Aldrich) with full anesthesia determined via toe pinch. DPOAE measurements were recorded using the TDT RZ6 processor and BioSigTZ software. The ER10B+ microphone system was inserted into the ear canal in way that allowed for the path to the tympanic membrane to be unobstructed. DPOAE measurements occurred at 8, 16, and 32 kHz with an f2/f1 ratio of 1.2. Tone 1 was *.909 of the center frequency and tone 2 was *1.09 of the center frequency. DPOAE data was recorded every 20.97 milliseconds and average 512 times at each intensity level and frequency. At each tested frequency, the stimulus intensity was reduced in 10 dB steps starting at 90 dB and ending at 10 dB. DPOAE threshold was determined by the presence an emission above the noise floor. Baseline DPOAE recordings were performed when mice were 7-8 weeks old and post experimental recordings occurred after 14 days following noise exposure. Threshold shifts were calculated by subtracting the pre-noise exposure recording from the post-noise exposure recording.

### Noise Exposure

Mice were placed in individual cages in a custom-made wire container. A System RZ6 (TDT) equipment produced the sound stimulus which was amplified using a 75-A power amplifier (Crown). A JBL speaker delivered the sound to the mice in the individual chambers. The sound pressure level was calibrated using an NSRT-mk3 (convergence instruments) microphone and all chambers were within 0.5 dB of each other to ensure equal noise exposure. Mice were exposed to 100 or 106 dB SPL noise for 2 hours with an octave band noise of 8 to 16 kHz.

### Tizaterkib Treatment

Tizaterkib (HY-111483) was purchased from MedChemExpress and administered to mice via oral gavage. Tizaterkib was dissolved in 10% DMSO (D8418, Sigma-Aldrich), 5% Tween 80 (9005-65-6, MP Biomedicals), 40% PEG 300 (192220010, ThermoFisher Scientific), and 45% 0.9% saline. Mice were either administered 25, 5, 0.5, or 0.1 mg/kg/bw. Mice were treated both morning and night (12 hours between treatments) for either 1, 2, or 3 total days. Treatment began 45 minutes (Figure 1) before noise exposure or 24 (Figures 2-5, 6-8) or 48 hours (Figure 6) after noise exposure. Mice were weighed periodically throughout the experimental protocol to monitor any drug toxicity and to ensure proper dosages were administered to the individual mice.

### Ctbp2 Staining and quantification

Mice were sacrificed following the post-experimental hearing tests and cochleae were dissected and placed into 4% PFA. Organs of Corti were micro dissected and co-stained with anti-Ctbp2 (1:800; 612044, BD Transduction) and myosin VI (1:400; 25-6791, Proteus Biosciences). Goat anti-rabbit Alexa Fluor 488 (1:400; A11034) and goat anti-mouse Alexa Fluor 647 (1:800; A32728) were purchased from Invitrogen as the secondary antibodies. Confocal Imaging was performed using a Zeiss 700 upright scanning confocal microscope with images taken with the 63x objective lens. Final images were achieved by taking a z stack image of the organ of Corti and processed through the ZenBlack program. The number of Ctbp2 puncta were counted per IHC with a total of 12-14 IHCs analyzed per region counted. The total amount of Ctbp2 puncta in each region were divided by the total amount of IHCs in that region to determine the number of Ctbp2 puncta per IHC.

### Cochlear Cryosectioning and CD45 Staining

FVB mice aged 6-8 weeks’ old were exposed to 100 dB SPL noise (8-16 kHz octave band) for 2 hours. Mice were either treated with carrier alone or 0.5 mg/kg Tizaterkib twice a day for 3 days beginning 24 hours after noise exposure. Mice were then sacrificed 1 hour after noise the last Tizaterkib treatment which was approximately 85 hours (little over 3 ½ days) after noise exposure. Another set of mice were sacrificed 8 days after noise exposure to observe a different time point. Cochleae were extracted from mice and placed in 4% PFA for 2-3 days. Cochleae were then decalcified in 120 nM EDTA for 2-3 days. Following decalcification, cochleae were transferred to a 30% sucrose solution and kept at 4°C overnight. The next day, samples were put into a solution of 30% sucrose and OCT compound (4583; Sakura) for 4 hours at 4°C. Samples were then placed in OCT compound overnight at 4°C. The next day, cochlear tissues were oriented in within cryomolds containing OCT compound and frozen on dry ice. Frozen tissues were cut with 10µm thickness. captured on glass slides, and allowed to dry for several hours.

Cochlear cryosections were then blocked and permeabilized in a solution of 5% FBS and 0.2% triton X-100 in PBS. Tissues were stained overnight at 4°C with mouse CD45 antibody (1:200; Af114, R&D Systems). The next day, tissues were then stained for one and a half hours with Alexa Fluor 568 donkey anti-goat (1:400; A11057, Invitrogen) and DAPI (1:1000; D1306, Invitrogen) to counterstain nuclei. Tissues were mounted in Fluoromount-G (00-4958-02, Invitrogen) and imaged using a Zeiss 700 upright confocal microscope. Post-acquisition images were analyzed using the IMARIS imaging software and automatically quantified following intensity thresholding. CD45 positive cells were then cross-checked manually to ensure positive CD45 cells were co-stained with DAPI. Raw CD45 cell counts were normalized in the scala tympani region to the area of the region counted and averaged as CD45 positive cells per µm^2^.

### Western Blotting

FVB mice aged 7-8 weeks’ old were exposed to 100 dB SPL noise (8-16 kHz) for 2 hours and treated with carrier or 0.5 mg/kg Tizaterkib twice a day for three days beginning 24 hours after noise exposure. Mice were sacrificed 6 days after noise exposure and whole cochlear lysates were prepared in lysis buffer (9803; Cell Signaling) after adding protease (cOmplete ULTRA Tablets 05892791001) and phosphatase (PhosSTOP 04906845001) inhibitors (Roche). Cochlea from each mouse were pooled together so each experimental group had tissues from 5 mice (10 cochlea). The lysates were centrifuged for 20 minutes at 16,000g at 4°C, and the supernatants were collected. Protein concentrations were determined with the BCA protein assay kit (23235, Thermo Fisher Scientific). 25 micrograms of total cell lysate were loaded on 10% SDS-polyacrylamide electrophoresis gel. After running the gel and transferring to a nitrocellulose membrane, the following antibodies were used for immunoblot analysis: anti-CD45 (1:1000; Af114, R&D Systems), anti-CD68 (1:1000; MCA1957, Bio-Rad), and anti-GAPDH (1:10,000; AB181602). Rabbit anti-goat (1:4000; 31402, Invitrogen), goat-anti rat (1:4000; 31470, Invitrogen), and goat-anti rabbit (1:4000; P0448, Dako) secondary antibodies were used. National Institute of Health (NIH) ImageJ software was used to quantify band intensities and recorded as the ratio to the GAPDH loading control. Each pooled cochlear lysate was run 3 times.

### Statistical Analysis

Statistics was performed using Prism (GraphPad Software). Two-way analysis of variance (ANOVA) or one-way ANOVA with Bonferroni post hoc test was used to determine mean difference and statistical significance.

## Author Contributions

T.T. and R.D.L. conceived the project. R.D.L. and M.A.I. performed the noise exposure and drug treatments in mice. R.D.L. and J.F. performed ABR and DPOAE tests. R.D.L., M.A.I., J.F., and T.T. analyzed ABR and DPOAE functional data. R.D.L. performed cochlear dissections and stained whole mount cochlear sections. R.D.L., M.A.I., A.T., and A.J. performed cochlear cryosections and stained images. R.D.L., M.A.I., A.T., and A.J. conducted confocal microscopy. R.D.L., A.T., and A.J. analyzed immunofluorescence data. R.D.L. performed and analyzed western blots. R.D.L. and T.T. wrote the manuscript with input from all authors. T.T. acquired funding for project and provided supervision.

## Acknowledgements

We thank Daniel F. Kresock, Kristina Ly, Dr. Christy Howe, Dr. Janee Gelineau-van Waes, Pat Steele, Ann Bryen and the Creighton University ARF staff for assistance with the mouse studies. We thank Dr. Robert Lewis from the Eppley Cancer Institute, University of Nebraska Medical Center, Omaha, NE for the kind gift of the KSR1 mice. The research was funded by the National Institutes of Health NIDCD grant 1R01DC018850 and American Hearing Research Foundation 2020 grant to T. Teitz. This investigation was conducted in facilities constructed with support from Research Facilities Improvement Program (G20 RR024001-01) from the National Center for Research Resources, NIH. The research was partially conducted at the Auditory and Vestibular Technology Core (AVT) at Creighton University, Omaha, NE (RRID:SCR_023866). This facility is supported by the Creighton University School of Medicine and grants GM103427 and GM139762 from the National Institute of General Medical Science (NIGMS), a component of the National Institutes of Health (NIH). IBIF was constructed with support from grants from the National Center for Research Resources (RR016469) and the NIGMS (GM103427). This investigation is solely the responsibility of the authors and does not necessarily represent the official views of the National Center for Research resources, NIGMS or NIH.

## The Paper Explained

### Problem

Hearing loss occurs in more than 10% of the world population with noise-induced hearing loss as one of the main causes, yet no FDA-approved drugs exist to prevent it. Hearing loss negatively impacts many aspects of individual’s daily life and hearing aids do not work for everyone; therefore, a treatment is desperately needed to prevent this highly common disorder.

### Results

We have shown that the drug tizaterkib, a highly-specific ERK1/2 inhibitor, protects mice from hearing loss when given orally before, and 24-48 hours after, moderate-to-high noise intensity levels. Protection was achieved with drug doses that are equivalent to the ones currently tested in humans for anticancer treatment, and no deleterious side effects were exhibited in the animals. The drug reduced nerve connectivity damage, and the mechanism of action was shown to be through the MAPK cellular pathway by taking advantage of a genetic knockout mouse model that has reduced activity of this specific pathway. Interestingly, we could show that while mice treated with noise alone had increased infiltrating immune cells in their cochleae for days after noise exposure, mice treated with tizaterkib and noise had reduced infiltrating immune cells in their cochleae to the low baselines levels measured in mice who were not exposed to noise.

### Impact

Our study provides evidence that targeting the MAPK pathway is a viable approach to mitigate noise-induced hearing loss. Tizaterkib is a promising preclinical compound that was shown to have a large therapeutic index in mice while offering up to 80% hearing protection. The immune response was tempered down with tizaterkib treatment, which supports regulating the immune response as a possible therapeutic strategy for reducing noise-induced hearing loss, and could be part of the mechanism by which ERK1/2 and MAPK inhibition confer hearing protection effects.

### For More Information

All data needed to evaluate the conclusions in the paper are present in the paper and/or Expanded View. Additional data related to this paper may be requested from the authors.

**<u>Expanded View Figure 1:</u> Representative ABR traces.** Representative post treatment ABR traces are shown from all different treatment groups shown in Figure 2. The threshold was recorded as the last trace with at least 3 of the 5 ABR waveforms present.

**<u>Expanded View Figure 2:</u> Tizaterkib treated mice have less CD45 positive cells in the stria vascularis compared to noise alone treated mice. (A)** Representative images of cochlear cryosections of the stria vascularis following noise and tizaterkib treatment that are stained with CD45 (red) and DAPI (blue). **(B)** Quantification of CD45 positive cells per experimental group. Carrier (grey), tizaterkib alone (green), noise alone (red), and noise + tizaterkib (blue). Data shown as means ± SEM, *P<0.05, compared to noise alone by one-way ANOVA with Bonferroni post hoc test. n=5-6

**<u>Expanded View Figure 3:</u> No positive immunostaining occurred with secondary alone and zoomed in images of CD45 positive cells. (A)** Representative images of cochlear cryosections stained with DAPI (blue) and 568 donkey anti-goat secondary antibody (red) (secondary by itself with no CD45 primary antibody). Top image is DAPI and secondary together and bottom image is secondary by itself. **(B)** Zoomed in images of CD45 positive cells. Top image is DAPI (blue) and CD45 (red) together and bottom image is CD45 by itself.

**<u>Expanded View Figure 4:</u> The number of CD45 positive cells are back to baseline levels 8 days following noise exposure. (A)** Representative images of cochlear cryosections of noise and noise + tizaterkib treated mice 8 days following noise exposure. Sections are stained with DAPI (blue) and CD45 (red). Image on top has DAPI and CD45 staining and image below has CD45 alone. **(B)** Zoomed in representative images of the scala tympani region 8 days after noise exposure. **(C)** Zoomed in representative images of the stria vascularis 8 days after noise exposure.

